# Dimensionality Reduction on Spatio-Temporal Maximum Entropy Models of Spiking Networks

**DOI:** 10.1101/278606

**Authors:** Rubén Herzog, María-José Escobar, Rodrigo Cofre, Adrián G. Palacios, Bruno Cessac

## Abstract

Maximum entropy models (MEM) have been widely used in the last 10 years to characterize the statistics of networks of spiking neurons. A major drawback of this approach is that the number of parameters used in the statistical model increases very fast with the network size, hindering its interpretation and fast computation. Here, we present a novel framework of dimensionality reduction for generalized MEM handling spatio-temporal correlations. This formalism is based on information geometry where a MEM is a point on a large-dimensional manifold. We exploit the geometrical properties of this manifold in order to find a projection on a lower dimensional space that best captures the high-order statistics. This allows us to define a quantitative criterion that we call the “degree of compressibility” of the neuronal code. A powerful aspect of this method is that *it does not require fitting the model*. Indeed, the matrix defining the metric of the manifold is computed directly via the data without parameters fitting. The method is first validated using synthetic data generated by a known statistics. We then analyze a MEM having more parameters than the underlying data statistics and show that our method detects the extra dimensions. We then test it on experimental retinal data. We record retinal ganglion cells (RGC) spiking data using multi-electrode arrays (MEA) under different visual stimuli: spontaneous activity, white noise stimulus, and natural scene. Using our method, we report a dimensionality reduction up to 50% for retinal data. As we show, this is quite a huge reduction compared to a randomly generated spike train, suggesting that the neuronal code, in these experiments, is highly compressible. This additionally shows that the dimensionality reduction depends on the stimuli statistics, supporting the idea that sensory networks adapt to stimuli statistics by modifying the level of redundancy.

**Author Summary:** Maximum entropy models (MEM) have been widely used to characterize the statistics of networks of spiking neurons. However, as the network size increases, the number of model parameters increases rapidly, hindering its interpretation and fast computation. Here, we propose a method to evaluate the dimensionality reduction of MEM, based on the geometrical properties of the manifold best capturing the network high-order statistics. Our method is validated with synthetic data using independent or correlated neural responses. Importantly, we show that dimensionality reduction depends on the stimuli statistics, supporting the idea that sensory networks adapt to stimuli statistics modifying the level of redundancy.

## Introduction

It is widely admitted that the spikes exchanged between neurons convey information encoded in a, yet unknown, manner. Deciphering these hidden “neural codes” - there is no reason why neurons should use a unique coding strategy - is a contemporary challenge. A natural approach consists of detecting statistical regularities in the spike patterns (“words”) produced by a population of neurons. This problem is challenging since the number of possible patterns grows exponentially with the number of neurons and the time window defining the words. A population of *N* neurons may produce 2^*N×τ*^ words of time length *τ*, but experimental recordings fortunately produce a quite smaller subset. In practice, once the number of neurons increases beyond 20, the total number of possible words becomes intractable. Yet, one has to extract, from the empirical statistics of the words displayed, a canonical model predicting the observed statistics, paving a possible way toward decoding. A theoretical framework to tackle this issue is based on the Maximum Entropy Principle (MEP) [42, 46]. The objective is to find the least structured model - the one with maximum entropy - given constraints provided by the average values of certain “features” or observables.

In its simplest form MEP restricts to spikes occurring at the same time. For example, the Ising model is constrained by the occurrence of spikes emitted at the same time by 2 distinct neurons (Model with “pairwise interactions”) [2, 3]. Most extensions with more neurons (triplets-quadruplets) restrict as well to spikes occurring at the same time [8, 9, 10]. The mathematical consequence is that, in these MEM, spikes events occurring at different times are independent. The probability of spike patterns occurrence at time *t* does not depend on the history. We call these models “MEMs without memory”. However, neurons interact together with some delay, inducing spatio-temporal correlations which should be considered as well as constraints for the MEM [8, 9, 10]. The MEP can be extended to handle these spatio-temporal correlations [12, 16] capturing and predicting the collective spatio-temporal pattern activity. This approach has been successfully used to characterize the spike train statistics in cortex cultures [11] and in the vertebrate retina network [33, 14].

A strong caveat of MEM is their number of parameters increasing rapidly with the size of the system. But one expects some redundancy in the spike activity of biological neuronal networks (in contrast e.g. with artificial neural networks with random uncorrelated synapses), reflecting a latent structure on the statistics of the activity. From this point of view, the dimensionality of the MEM (the number of its parameters) could be reduced. There might be some analogy with signal processing compression, where the presence of statistical redundancy can be exploited to map the signal onto a subset of independent channels (i.e. non-redundant). As shown in [23, 13], the reduction is more pronounced if the parameters are related by hidden mathematical dependencies.

Several tools are available to reduce the dimensionality of a given statistical model, based on the trade-off between dimensionality and likelihood, e.g Akaike Information Criterion (AIC), Bayesian Information Criterion (BIC) or Minimum Description length (MDL) among others. However, these methods necessarily need to fit different models to the same dataset to find the optimal dimension. This can be prohibitive in the case of MEM when considering a large number of neurons. In contrast, a formal framework to find the optimal dimensionality of MEM based on the geometrical properties of the manifold of probability distributions -where a MEM is a point in large dimensional space, whose coordinates are the parameters-was developed in [41]. It generalizes over standard methods (AIC and BIC), taking into account simultaneously the dimensionality, the likelihood and the geometry of the manifold formed by the family of statistical models. Here, we propose to exploit the geometric structure of the manifold formed by the MEM’s to find an optimal set of dimensions capturing the information contained on neural spiking data. Our approach generalizes previous approaches considering only i.i.d samples [41], and is general enough to consider spatio-temporal interactions at several lags between neurons. It relies on the spectral properties of a matrix, called “Fisher metric” in statistics and information geometry [24] and “susceptibility Matrix” in statistical physics, capturing the effect of MEM parameters variations on the second-order statistics of the network activity. Remarkably, as we show here this matrix can be numerically computed from data without fitting the MEM parameters, allowing us to apply the method to a large set of neurons (*∼*100 neurons). Based on the spectral properties of this matrix, which reflects the local geometry of manifold, we determine the optimal model dimensionality that still preserves the correlation structure in neural activity.

The method is first validated using synthetic data generated with a known underlying statistics. It is then applied to the spiking response in a neural population of retinal ganglion cells (RGC) *in vitro*, recorded from a diurnal rodent Octodon degus [26] using a 252-MEA (multi-electrode array). We consider three sets of neural responses obtained after applying three type of visual stimuli: i) spontaneous photopic activity; ii) white-noise checkerboard and iii) a short natural movie. From the mathematical analysis and from an analogy with signal compression, we define a “compressibility” criterion for the MEM. Based on our experimental observations, we found that RGC activity is highly compressible (∼ 50% of the imposed model dimensionality) and that the stimuli spatio-temporal modulation increases the number of independent dimensions required to optimally represent the neural activity. This suggests that RGC population activity adapts to stimuli conditions, changing the number of “coding channels” according to stimuli correlations.

## Methods

### Spike trains

We consider the spiking activity data from a population of *N* interacting neurons. This data usually comes from experimental recordings using multi-electrode-arrays in the retina or cortical areas. We assume that there is a minimal time interval such that any neuron fires at most one spike within this interval. This provides a time discretization usually referred as “binning”. The *spike-state ω*_*i*_(*n*), takes the value 1 whenever the *i*-th neuron emits a spike at the time bin *n*, otherwise is zero. The *piking pattern* 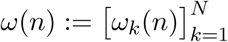 is the spike-state of the entire network at time *n*. The *spike block* 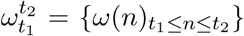 represents the activity of the whole network between time bins *t*_1_ and *t*_2_. The *length* of a spike block is the number of time steps *t*_2_ − *t*_1_ +1. Experimental data consist of a *spike-block* of finite size *T*, denoted by 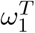. In this paper we also consider infinite spike sequences 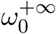 denoted by *ω* to alleviate notations. The state space of an infinite sequence of spiking patterns is denoted by Ω.

### Observables and Monomials

We call *observable* a function *f*: Ω → ℝ, that associates a real number to a spike-train. We say that *f* has range *R*, if for every pair of spike trains *ω, ω′ ∈* Ω we have that *f* (*ω*) = *f* (*ω′*) if and only if 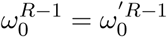, that is, *f* only depends on the first *R* spike patterns of the spike-train. An important class of observables are the *monomials*, which are binary observables consisting of finite products of spike states, given by:

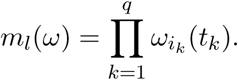

If one fixes a finite set of pairs 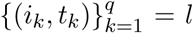(neuron index, time index), there are finitely many such possible monomials, which can be indexed by an index *l* in one-to-one correspondence with the set of pairs (*i*_*k*_, *t*_*k*_). The observable *m*_*l*_(*ω*) = 1, if and only if neuron *i*_*k*_ spikes at time *t*_*k*_, *∀k ∈* {1, *…, q*} in the spike-train *ω*, where *q* is the number of spike states in the observable, and *m*_*l*_(*ω*) = 0 otherwise. In a range *R ≥* 1 monomial, the firing times *t*_*k*_ are constrained within the interval {0, *…, R* − 1}.

### Inference of the spike train statistics via Maximum Entropy principle

A generalized version of the maximum entropy approach can be framed rigorously using the ther-modynamic formalism of subshifts of finite type [38], which offers a way to build maximum entropy Markov chains (MEMC) from data in a principled way through a variational principle [16, 13]. As we will see, framing the maximum entropy problem in this way is particularly useful, as set up the conceptual framework to exploit ideas from thermodynamics and information geometry.

We assume that the spiking data is a sample of a time homogeneous Markov chain of memory *R*, i.e., taking values in 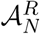 in other words, 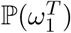can be decomposed according to:

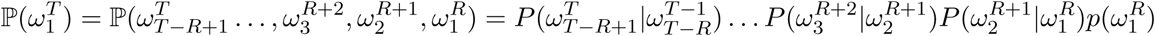

where 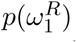 is the probability of the initial state and 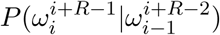 is the conditional probability or the transition matrix elements. The goal of the inference problem is to estimate the transition matrix *P*.

### Entropy Rate

If the spike train *ω* is characterized by a stationary ergodic Markov measure denoted by *μ*(*p, P*) taking values in 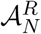 with homogeneous transition matrix *P* and unique stationary distribution *p*, the entropy rate of *ω* is referred as *Kolmogorov-Sinai entropy (KSE)* of *μ*(*p, P*) and takes the simple form [39]:

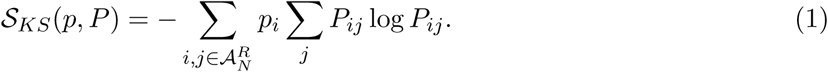

where *P*_*ij*_ = *P* (*j|i*). It is easy to see that when the stochastic process is i.i.d (*P*_*ij*_ = *p*_*j*_) we recover the classical definition of Shannon entropy.

### Variational Principle

We now introduce the maximum entropy principle (MEP) in the context of Markov chains (it extends to chain with infinite memory). In comparison to the standard statistical physics formulation of MEP, the present formalism extends to time-dependent interactions including with an infinite range (requiring then appropriate summability conditions [37].

Given a set of observables (monomials) *m*_*l*_, *l* = 1 *… L*, we denote *ν*[*m*_*l*_] the expectation of *m*_*l*_ under the probability *ν*. We also denote *c*_*l*_ the empirical average of *m*_*l*_ (i.e. measured from an experimental raster). The MEP find the unique invariant probability *μ* satisfying *μ*[*m*_*l*_] = *c*_*l*_, *l* = 1 *… L* that maximize the KSE. This is equivalent to solve the following problem considering observables of range *R ≥* 2:

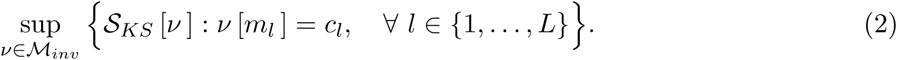

where ℳ_*inv*_ is the set of translational invariant measures (stationary). Since the function *ν → S*_*KS*_ [*ν*] is strictly concave, there is a unique maximizing Markov measure *μ*(*p, P*) given the constraints *c*_*l*_. This probability is uniquely defined by the *potential*:

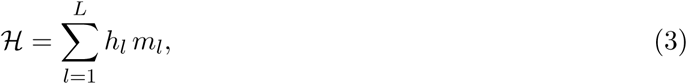

This is a linear combination of the monomials *m*_*l*_ associated with constraints. The coefficients *h*_*l*_ are called the parameters of the model as there variation change the probability of events. They also formally correspond to interactions between neurons (e.g.; in the Ising model *h*_*l*_ are either the pairwise interaction *J*_*ij*_ between two neurons - monomials of type *ω*_*i*_(0)*ω*_*j*_(0) - or the local external magnetic field - monomials of type *ω*_*i*_(0)).

Equation (2) is equivalent to the following unconstrained problem, which is a particular case of the so-called *variational principle* of the thermodynamic formalism [38]:

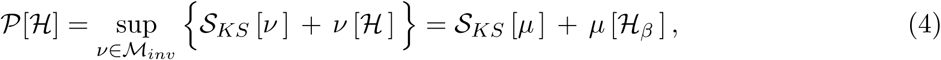

where 𝒫 [ℋ] is called the *free energy of ℋ*, and 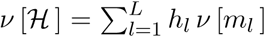 is the average value of ℋ with respect to *ν*.

The average value of the observables, their correlations, as well as their higher cumulants can be obtained by taking the successive derivatives of the free energy with respect to the parameters *h*. This outlines the important role played by the free energy in this framework. In particular, taking the first derivative:

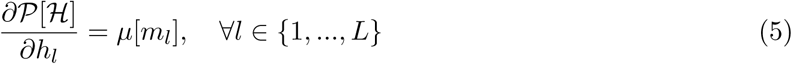

where *μ*[*m*_*l*_] is the average of *m*_*l*_ with respect to *μ*. The second derivative of the free energy w.r.t a single parameter *h*_*l*_ gives the second cumulant or the variance of the observable *m*_*l*_:

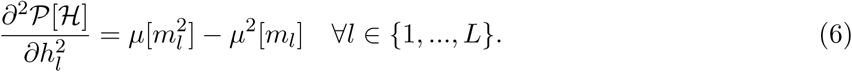

### Susceptibility matrix

Let *φ* be the shift map *φ*: *ω → ω*, defined by 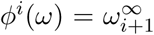Let *m*_*l*_ be an arbitrary observable. We may consider the sequence *{m*_*l*_ *º φ*^*i*^(*ω*) as a random variable whose statistical properties depend on those of the process producing the samples of *ω*. For a pair *m*_*l*_, *m*_*l*_′ of monomials, the *time covariance of order r* of the stationary processes *{m*_*l*_ *º φ*^*n*^; *n ≥* 0} and *{m*_*l*_*t º φ*^*n*^; *n ≥* 0} is defined as:

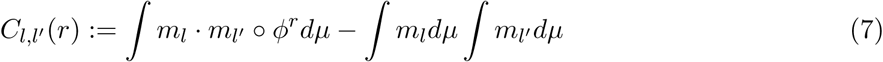

In particular, the auto-covariance of order *r* is:

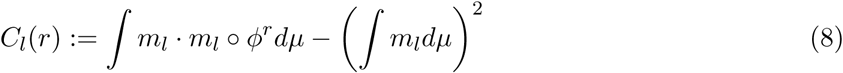

For a pair of finite range observables *m*_*l*_, *m*_*l*_′, the susceptibility can be obtained from the free energy as follows:

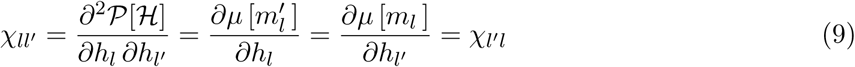

It is also a standard result in ergodic theory (Green-Kubo formula) that the elements of the matrix *χ* can be obtained via time correlations:

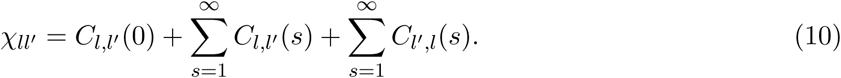

Especially, for the diagonal term:

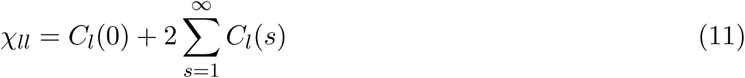

**Remark 1:** The matrix *χ* is at the core of our analysis and can be computed directly from data, assuming that *μ* is the empirical measure, that is without fitting the maximum entropy parameters.

**Remark 2:** In the case of memory independent MEM (e.g. Ising), and *only in this case*, (9) reduces to *χ*_*ll′*_ = *C*_*ll′*_ (0). In general, *χ* involves correlations at all times

### Properties of the susceptibility matrix

Our method relies on the analysis of this matrix, which has the following properties:

i. The set of all parameters fixes the statistical model. A slight variation *δh*_*l*_ of the sole weight *h*_*l*_ affects all the other monomials average. One can show that 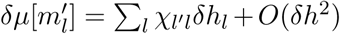. Thus, *χ* is a linear response matrix.
ii. It is symmetric and positive, thus with real positive eigenvalues, which can be arranged increasingly *λ*_1_ *≥ λ*_2_ *≥ … λ*_*k*_ *≥ λ*_*L*_ *>* 0.
iii. The set of 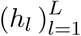 is a vector of *L* dimensions, which can be seen as a point in 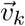The set of eigenvectors ***v***_*k*_ of *χ* constitute an orthogonal basis of this space, where the *k*-th eigenvector *_v*_*k*_ is the *k*-th direction in the space ε.
iv. The closest two points are in ε, the closest are the statistics they predict. As a corollary, trying to fit empirical statistics, two models corresponding to “close” points might be indistinguishable as they describe equally well the empirical statistics. The notion of closeness is however ambiguous here and requires to define a proper metric. This is precisely what *χ* does: it defines a metric (called Fisher metric).
v. From a statistical perspective,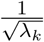 gives the amplitude of second-order fluctuations in the estimation of coefficients *h*_*l*_ projected on direction 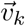. The smaller the eigenvalue, the larger the amplitude. Finally, from the linear response perspective, 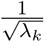 tells us how much a small variation in the estimation of average of monomials affects the estimation of *h*_*l*_.

The Susceptibility matrix (or Fisher matrix) is used here in order to study *both* the geometric structure and the statistical fluctuations of the family of statistical models [24, 36, 23].

### Distinguishable MEMC

Consider a spike train data set 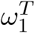 consisting of *T* spike patterns generated by a biological neuronal network. Given a set of *L* arbitrary observables 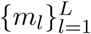 (possibly non-synchronous) we compute their empirical averages from 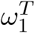 in order to set the constraints and infer the maximum entropy parameters ℋ = (*h*_1_, *…, h*_*L*_) characterizing the MEMC. We may use these parameters to generate a sample 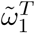 of the inferred MEMC of same size *T* as the original data set. Considering the same set of observables we can apply again the MEP to infer a new set of parameters ***h′*** from 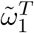, which is expected to be different from ℋ due to finite size sampling. Thus, the MEMC specified by *H***_*h*_** and *ℋ*_*h′*_ cannot be distinguished on the basis of a dataset of finite length. However, increasing the sample size, one expects the MEMC specified by the potential *ℋ*_*h′*_ to get “closer” to the one characterized by *H***_*h*_**. This idea can be rigorously formulated using large deviations techniques (see appendix).

**Definition** Consider two MEMC specified by *H***_*h*_** and *ℋ*_*h′*_ within the same family of observables. Then, given a dataset of length *T* and empirical averages sampled from the model specified by **_*h*_** and tolerance ϵ *>* 0, we say that the MEMC with parameters ℋ and ***h′*** are *E*-*indistinguishable* if:

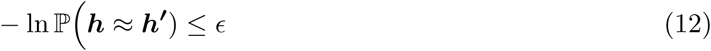

The notation *≈* stands for “inside a ball of radius *δ*, for some small *δ >* 0, as explained in the supplementary material. This last property identifies an approximatively elliptical region of *E*-indistinguishable models around each MEMC specified by ***h***, whose volume can be easily calculated in the large *T* limit [41, 36].

### Volume of indistinguishable models

Following Balasubramanian [41] two distributions *μ*^(1)^, *μ*^(2)^ are indistinguishable with tolerance ϵ if *-* ln 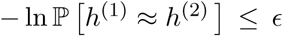If *T* is large enough, the set of indistinguishable distributions defines an ellipsoid with a volume denoted by 𝒱 called *confidence volume*. Points inside the confidence volume correspond to indistinguishable models within a tolerance.ϵ When the sample size *T*, the model dimension *L* (number of observables-parameters) and the tolerance *ϵ* = −log *κ* are fixed the volume is:

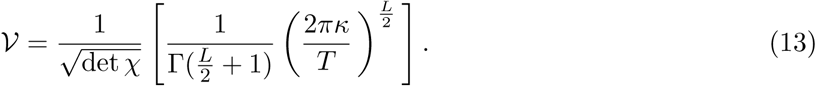

Therefore 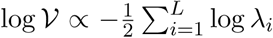. We observe that the model estimation is better if the eigenvalues of *χ* are larger, which implies that there exists a set of eigenvalues that can be neglected given their small magnitude, i.e. huge fluctuations impairing the model estimation. Then, instead of considering the volume in the space of all parameters, let us consider the volume 𝒱(*k*) of the projection in the subspace spanned by the *k* first eigenvectors of *χ*. We have:

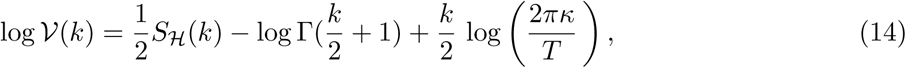

with:

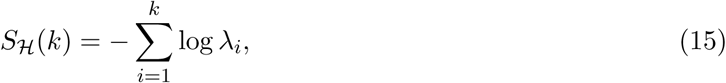

where the eigenvalues *λ*_*i*_ are ordered decreasingly. In eq. (14), the second term, 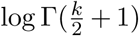, depends only of *k*. The third term, 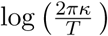 characterizes the effect of accuracy and finite sampling. Therefore, the only term which depends on the statistical model (here characterized by the potential ℋ) is *S* _*ℋ*_ (*k*). In particular, it depends on the number of neurons *N* and the range *R* of the potential.

As stated above, the inverse of the eigenvalues tells us how much a small variation in the estimation of the parameters affects the statistics. For a large eigenvalue *λ*_*k*_, a tiny variation in the direction *v*_*k*_ has a dramatic impact on the general statistics. On the opposite, small eigenvalues correspond to sloppy dimension where even a big change in the corresponding direction has small impact. The notion of stiff and sloppy dimension in statistics is not new and has been used by several authors, including for the analysis of spike trains [25], but the treatment we propose, for MEM with spatio-temporal interactions (in contrast to previous papers dealing with Ising model) is, to our best knowledge, a novelty. Thus, using *χ* as a metric to distinguish between models, we can find the optimal number of dimensions of a MEM given data, which is the main goal in this paper.

### Dimensionality reduction

In information geometry, one extends a family of parametric probability distributions to a manifold ℳ such that the points in ℳ are in a one to one relation with the probability distributions. The parameters of the distributions can thus also be used as coordinates on ℳ.

For a fixed potential form (3),the MEMC parametrized by the coefficients ***h***, corresponds to a point in the space ε. Equivalently, this point corresponds to a unique Markov measure. If the *h*_*l*_s are tuned independently from each others, the set of MEMC fully spans ε. However, when fitting data from neuronal networks, either artificial or biological, one expects hidden relations between the *h*_*l*_s, constrained by dynamics [13]. In this case, the MEMC spans a manifold ℳ in ε, of (presumably) quite lower dimension.

To estimate the dimension of ℳ, i.e, the dimension of the manifold of ε sufficient to explain the data with a minimal redundancy we use the following argument. Considering the equation (15) as the eigenvalues *λ*_*k*_ increases, there will be one or more *k*s at which *S* _*ℋ*_ (*k*) is expected to become bigger than the sum of the other two terms of (14), which are both negative. This means that, for increasing *k*, log 𝒱(*k*) will first decrease then it will increase, yielding at least one minima on the function. There is therefore a critical value of *k*, denoted by *k*_*c*_ *≡ k*_*c*_(ϵ, *T*), which characterizes the optimal dimension for which the *volume is minimal*. The value of ϵ belongs to some interval; outside this interval log 𝒱(*k*) is convex or concave having only trivial minima. Inside this interval, we obtain a set of *k*_*c*_ *≡ k*_*c*_(*T*) minimizing the volume, which characterize the number of dimensions ensuring that the model indeterminacy is minimized (see figure 1). Additionally, *k*_*c*_ provides a rough estimate of the dimension of ℳ.

**Figure 1:**
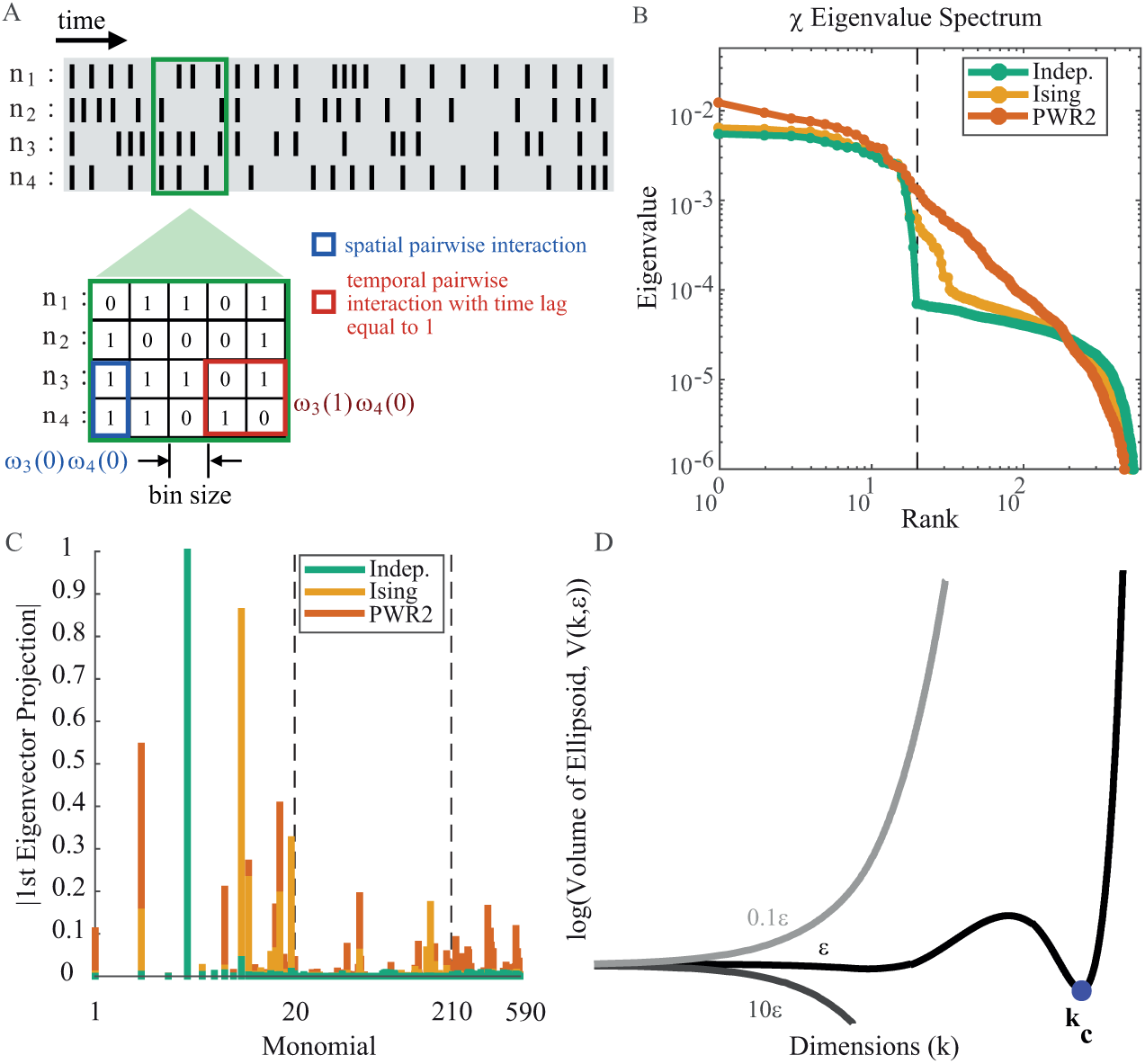
Dimensionality reduction framework overview. A: Raster representing the binary activity of *N* = 4 neurons (rows) over time (*T*). Green square shows a slice of this raster. For this raster 2 types of pairwise interactions are defined: spatial interactions (blue) and temporal interactions with *R* = 2, i.e. one time-step between spikes (red). Spatial interaction is exemplified as *ω*_3_(0)*ω*_4_(0) (both neurons firing at the same time) and *ω*_3_(1)*ω*_4_(0) (neuron 3 firing one bin after neuron 4). B: Susceptibility matrix eigenvalue spectrum in log-log scale for Independent (green), Ising (orange) and Pairwise Range=2 (PWR2, red). Black vertical dashed line denotes the network size (*N* = 20). Indep. raster has a sharp cut-off close to the network size; Ising has a cut-off few eigenvalues beyond the network size (*k*_*c*_ = 29); PWR2 shows a monotonic decay of the eigenvalues magnitude, without a clear cut-off. C: First eigenvector absolute projections for Independent (green), Ising (orange) and Pairwise Range=2 (PWR2, red) rasters. Left-most vertical dashed line is the limit between spike rates and spatial interactions monomials while the right-most is the limit between spatial and temporal interactions monomials. Indep. raster projects only on rates monomials, while Ising projects on rates and spatial interactions, but not ont temporal ones. PWR2 projects on the tree types of monomials. D: Log of the volume of indistinguishable models, log 𝒱(*k, ϵ*) as the degrees of freedom (*k*) increases. The volume reaches a minimum (blue dot) for *k* = *k*_*c*_. Increasing *k* beyond *k*_*c*_ makes the volume explode. Increasing/decreasing the error (ϵ) one order of magnitude yields to strictly increasing or decreasing functions, where volume minimization is trivial and uninformative.

### General Method Description and Applications to Synthetic Data

In order to test the method in a context where the ground truth is known, we artificially generated spike trains from MEMs using different sets of observables:

1. Independent model: This model considers that neurons are independent, i.e. it only contains self-interactions (firing rates). The parameters of the potential control the firing rate of each neuron. A rate model with *N* neurons has therefore *L* = *N* parameters. We call this model Indep. to alleviate notations. The potential for this model reads:

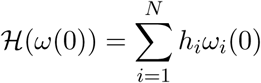
2. Ising model: This model considers firing rates and pairs of neurons firing at the same time (spatial). There is no interaction between spikes at different times. Considering *N* neurons, the model has 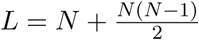 parameters. The potential for this model reads:

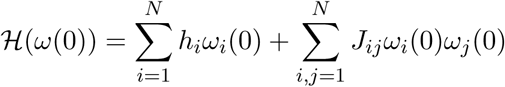
3. Spatio-temporal pairwise interactions model with memory depth 1: This model considers firing rates, pairs of neurons firing at the same time (spatial) and also pairs of neurons firing with one time-step delay between them (temporal). We call this model PWR2. Considering *N* neurons, this model has 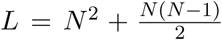 parameters (temporal self-interactions are not considered). The potential for this model reads:

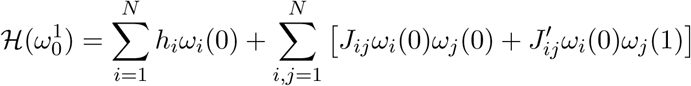 Note that, in contrast to *J*_*ij*_*s*, 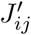 are not symmetric.
4. Scaled PWR2: We multiply the potential ℋ by a factor (corresponding to an “inverse temperature”). In our experiments we use *β* = {0.4, 0.6, 0.8, 1.2, 1.4}.

## Results

We apply our method to study the critical dimension (*k*_*c*_) of synthetic spike trains of size (*N* = 20, *T* = 10^6^) generated from random potentials corresponding to the Independent, Ising and PWR2 models. We generate 100 different rasters of each using the same set of parameters for each MEM. For each spike train, we compute the *χ* matrix *using the observables of the PWR2 model*, that is, we over fit when data are generated by an Independent or an Ising and, then, obtain the respective eigenvalue spectra.

We show in Fig 1B, the spectrum of *χ*. The entries of this matrix are estimated from data considering three different cases. First data was generated by a PWR2 model (red curve); in the second case, data comes from an Ising model (orange); in the last case data comes from an Independent model (green). In all cases, the dimension of *χ* is the same, but, in the independent and Ising case, we are overfitting the estimation, as for the independent model firing rates are enough to fit the data, and for the Ising model rates and pairwise interactions are enough.

The difference in the three cases is clearly seen in the spectrum of *χ*. Moving along the spectrum from left to right (increasing index *k*, decreasing the magnitude of the eigenvalue *λ*_*k*_) we observe a first sharp decrease (*cut-off*) at *k* = *N*, for the Indep. (Fig 1B, black) and Ising rasters.

In addition to the differences on the eigenvalue spectrum, the functional relationships between the monomials change depending on the underlying statistics, as exemplified by the first eigenvector of the corresponding *χ* matrix for each raster (Fig 1C). The Independent model shows large projections only on the monomials related to spike rates, while showing negligible projections on the pairwise monomials. The Ising model shows large projections as well on the spike rates monomials but in addition shows some projections on the spatial interactions monomials. Finally, the PWR2 raster has projections on all the monomials, reflecting in part its underlying statistics. Thus, the differences on the energy function are captured both by the eigenvalues spectrum and also by the structure of the corresponding eigenvectors.

In order to illustrate the volume of indistinguishable MEM, 𝒱(*k, ϵ*), we computed it at different ϵ and *k* finding some functions with non trivial minima, suggesting a reducible MEM (i.e. *k*_*c*_ *< L*). The presence of a cut-off on the eigenvalue spectrum shows that there is a value of *k* where the confidence volume is minimal (Fig 1D), i.e., where the model is more accurately determined. This point is highly non trivial and captures somewhat the role of spatio-temporal interactions on second order statistics^1^.

Finally, *k*_*c*_ is conditioned to the tolerance value ϵ as shown on Fig 1D. Increasing or decreasing ϵ one order of magnitude yields functions with trivial minima. This constrains the search of the minima to a subset of ϵ values. Extending the results for different values of ϵ shows that the number of dimensions related to the minimal volume decays monotonically with the accuracy (*κ* = *-* log *E*) until it reaches an inflection point; this inflection point is the *k*_*c*_ that we use as the number of relevant dimensions (Fig. 2).

**Figure 2:**
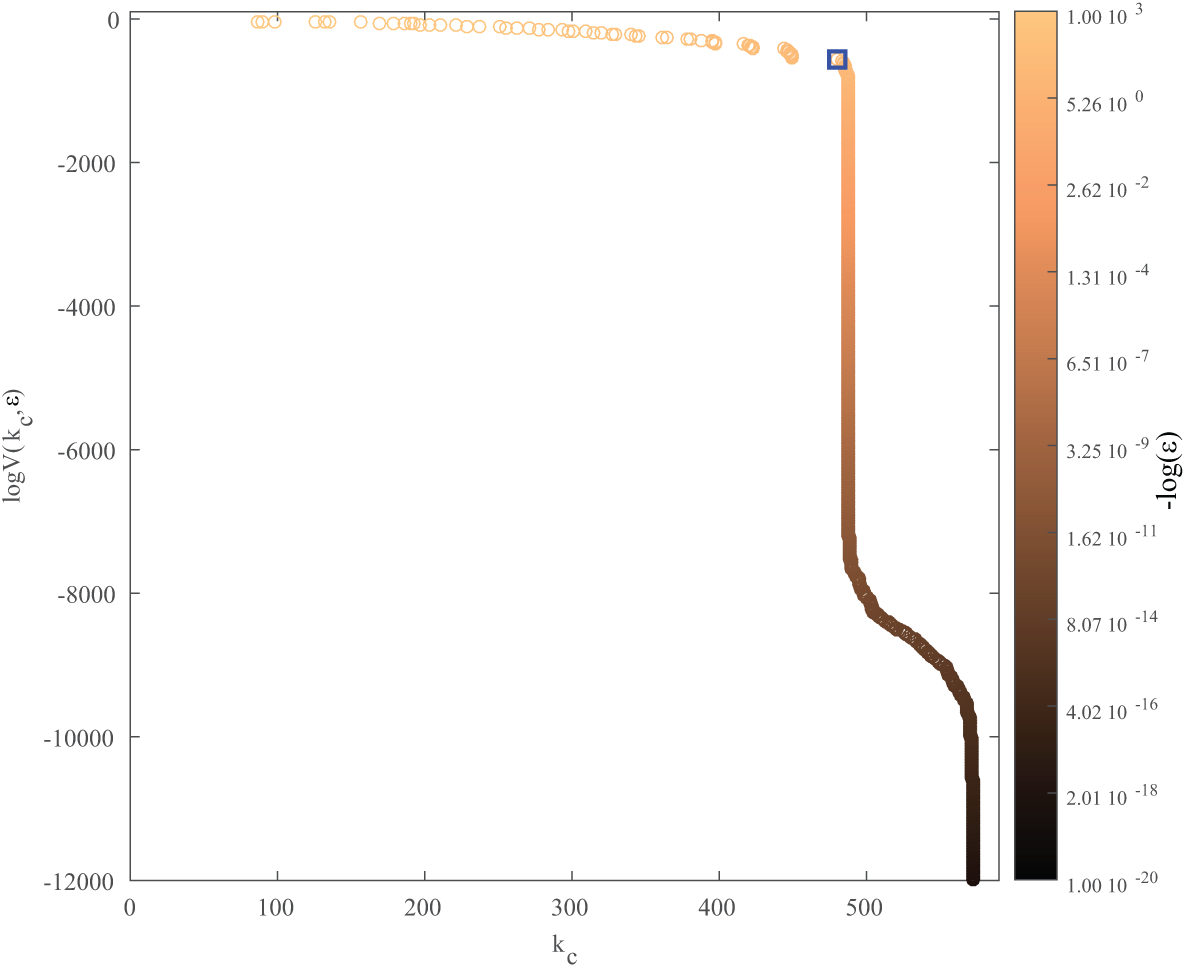
Finding *k*_*c*_ on the set of convex volume functions. The volume at the minimum and its corresponding *k*_*c*_ and *κ* (−log *E*) values. There are many convex functions and corresponding *k*_*c*_ that we could use, but given that we are looking for a cut-off, the actual *k*_*c*_ used is the last *k*_*c*_ before the inflection point (blue square) on the log *V*(*k*_*c*_, *E*) vs *k*_*c*_ curve, representing a trade-off between maximal number of dimensions, the highest accuracy and the minimal volume as possible. This is the method used to choose the *k*_*c*_ for all the data analyzed.

### *k*_*c*_ values on the independent and spatio-temporal correlated cases

Using the synthetic data, we evaluated how the optimal dimension given by *k*_*c*_ depends on the underlying statistics of neural data. Considering the *k* biggest eigenvalues, we computed the log of the volume *V*(*k, E*) (14) and compute its minimal value obtaining *k*_*c*_ (Fig 3A). The values of the parameters of the underlying MEM related to firing rates and to pairwise interactions were randomly chosen from a normal distribution with mean −5 and −1, respectively, and 1 standard deviation. We took one set of parameter for a PWR2 MEM and to obtain the Independent and Ising rasters, we kept only the parameters related to firing rates in the former and the ones relates to firing rates and spatial interaction in the latter. For this example we observe the following:

**Figure 3:**
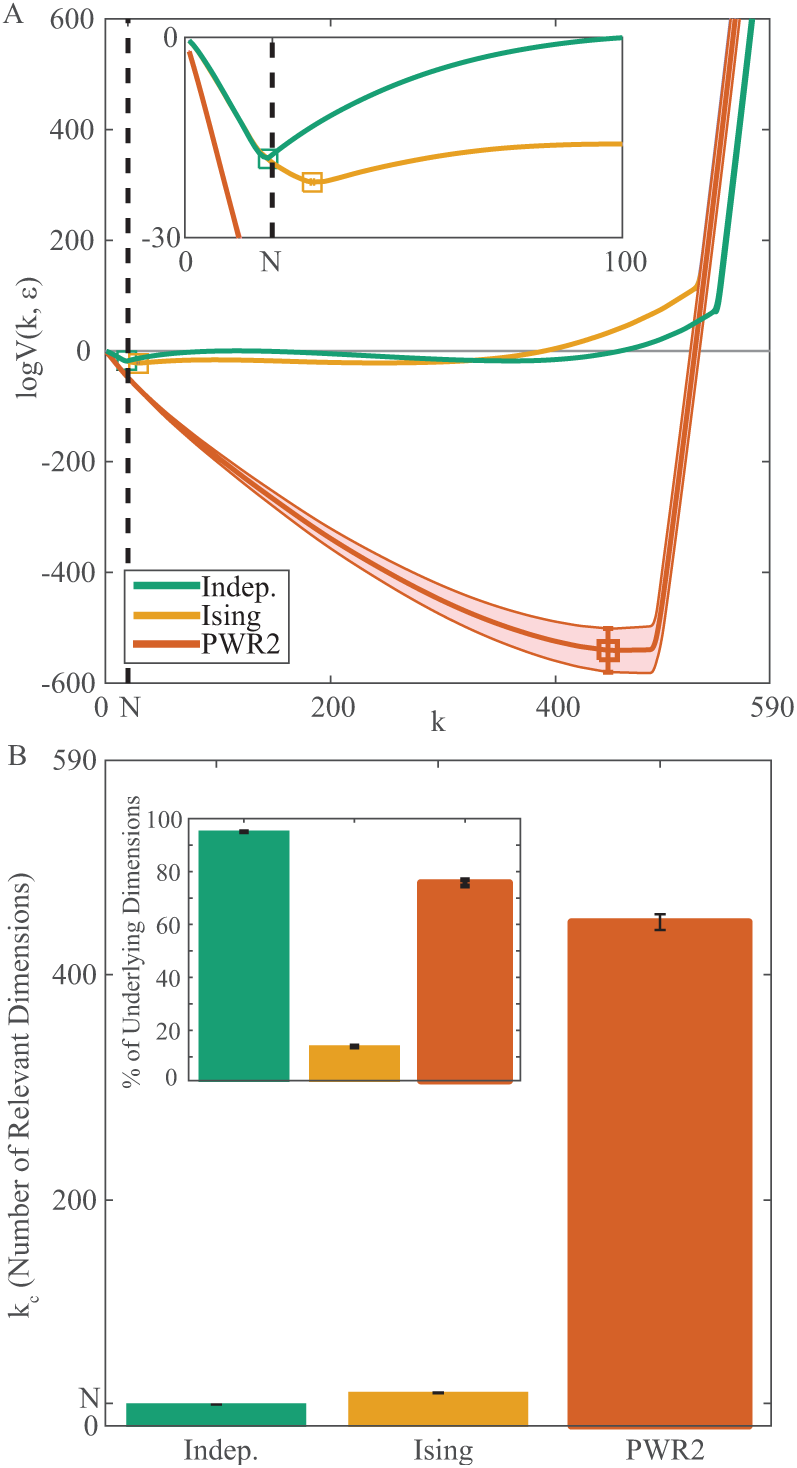
Dimensionality reduction by minimization of the volume of indistinguishable models. A: log of volume of indistinguishable models (log *V*) as function of the number of dimensions (*k*) at the maximal accuracy that yields no trivial minima (*κ* = 1336.6 *±* 16.00 for Indep., 1250.6 *±* 101.86 for Ising and 108.1*±* for PWR2). Thick solid lines are averages, shaded area is *±*1 s.d. of the 100 samples and black vertical dashed line is the network size (*N* = 20). The minimum of this function (squares) is the optimal number of dimensions capturing the raster statistics at the given accuracy, i.e. *k*_*c*_. Inset shows a zoom-in on the first 100 dimensions, focusing on the Indep and Ising. minima. B: Summary of the number of dimensions for both statistical models. We found a close relationship between the underlying model dimensionality and the number of dimensions required for an optimal model. In this example, *k*_*c*_ is 19 *±* 0 for Indep., 29.16 *±* 0.37, and 447 *±* 7.00 (mean *±* std) for Indep. and PWR2, respectively. Inset: *k*_*c*_ as a percentage of the underlying model dimensionality. Indep case show almost 100%, while Ising has *±* 14% of reduction and PWR2 *±* 75%.

In the case of independent statistics, the value of *k*_*c*_ is closely related to the network size (*k*_*c*_ =19 ± 0). In this example, one neuron has a very low firing rate (∼ 10^*-*5^) compared to the others (*>* 10^*-*4^). This explain the sharp cut-off at this neuron value. On the opposite, in the case of PWR2 the number of optimal dimension is *k*_*c*_ = 447 *±* 7.00 (see Fig 3B). The intermediate case, Ising, shows few number of dimensions more than *N*, but much less than *L*(590).

Interestingly, even when the number of dimension for the PWR2 and Ising is larger than the Independent case, the percentages of the optimal number of dimension relative to the total number of dimensions is smaller for them, compared to Indep. as shown in Fig 3C. This shows that our method can detect how many dimensions are needed for a MEM to fit data respect to underlying raster statistics.

For the Indep. case we found that *k*_*c*_ corresponds almost to the full dimensionality of the underlying model (*L* = 20), while Ising can be highly reduced respect to its full dimensionality. For PWR2 *k*_*c*_ corresponds to approximately 75% of the full underlying model dimensionality (Fig 3CB, inset), possibly related to finite size sample effects (events that are unlikely to be observed in finite time). So, for these examples, the effective dimensionality (*k*_*c*_) found by this dimensionality framework increases with the number of terms of the underlying statistics defined by the energy function.

### *k*_*c*_ depending on the the network size and recording length

We evaluated the impact of the number of neurons *N* and the raster duration *T* on the estimation of *k*_*c*_. We focus only on the Independent and PWR2 case, leaving Ising outside this analysis.

Fixing the neural population size (*N* = 20), we first varied the recording length from *T* = 10^2^ up to *T* = 2 *·* 10^6^ bins and observed the effect obtained (see Fig 4A) on the *χ* spectrum for the Independent (left) and PWR2 (right) cases. In the independent case, the first cut-off of the spectrum is increased while the value of *T* increases. On the opposite, in the case of PWR2 the cut-off is not observed, indeed, the separation in the spectrum of dimensions related to neuron firing rates or combined activation is less evident than the Indep. case. Additionally, we obtained the *k*_*c*_ value for each value of *T* as shown in Fig 4B. In both cases, we can consider that the estimation of *k*_*c*_ converges for values of *T* over 10^6^ bins.

**Figure 4:**
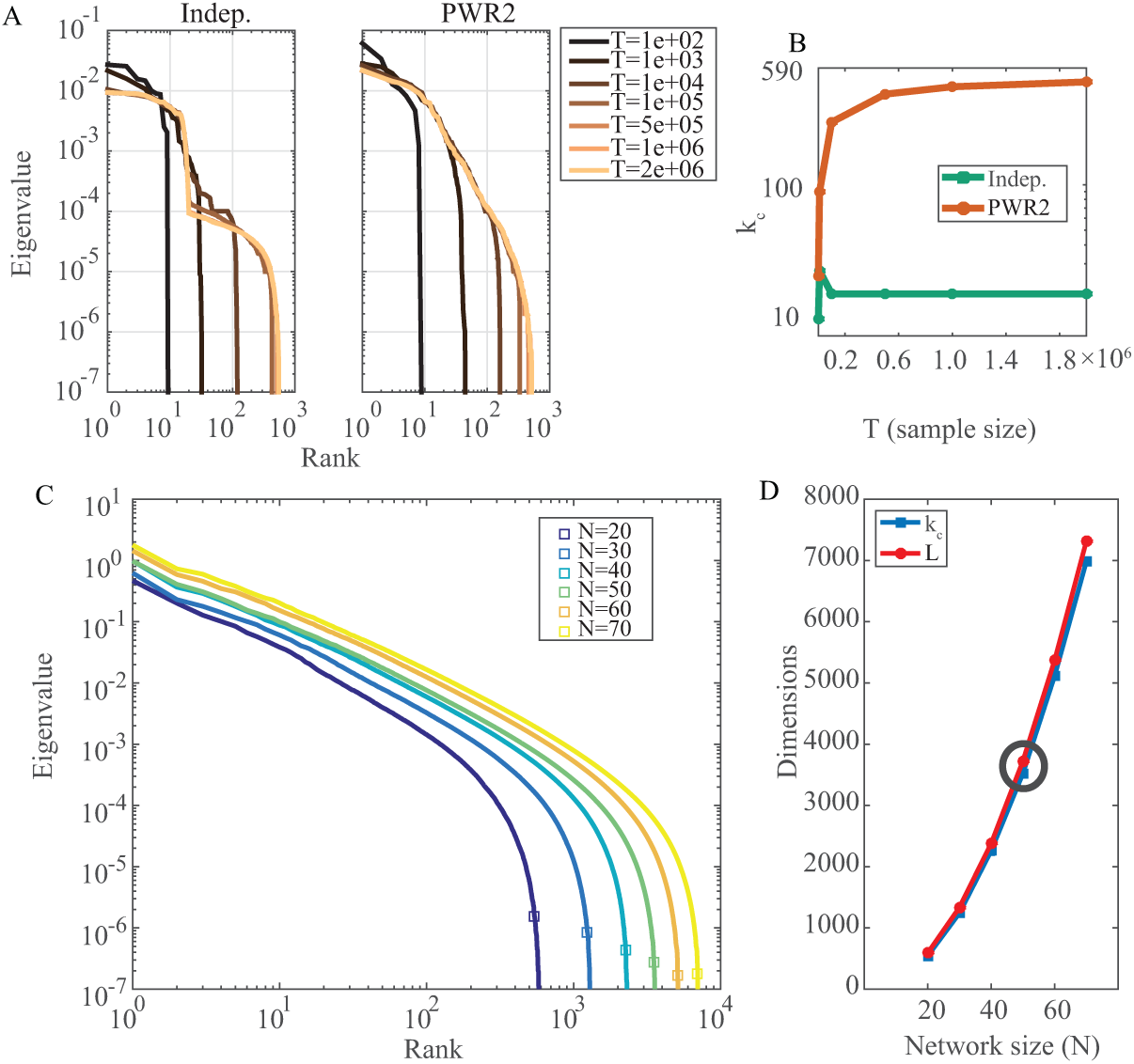
Method dependence on recording length and network size. A: Spectra of Indep. and PWR2 rasters using different recording lengths (colors) for *N* = 20. B: *k*_*c*_ values for both rasters at different recording length. Using *T ≥* 10^6^ is enough to have good *k*_*c*_ estimations. C: Spectra for random PWR2 rasters with increasing network size (*N*, colors) with the corresponding *k*_*c*_ on it (squares). The length of the spectrum (number of eigenvalues) changes because of the increase of model parameters, but the shape of the spectrum remains the same given the constant underlying model. D: *k*_*c*_ value for each network size (blue) and corresponding maximal dimensionality (*L*, red). Black circle denotes the raster size that we use in biological recordings (*N* = 50). *k*_*c*_ *∼ L* at almost all network sizes, but as *N* increases, the number of possible interactions grows, requiring longer recording lengths, making *k*_*c*_ diverge from *L*.

Fixing now the value of *T* = 10^6^, we evaluated the effect of the network size *N* on the estimation of *k*_*c*_ for the Independent and PWR2 case. As shown in Fig 4C, increasing the number of neurons increases the model dimensionality as well as *k*_*c*_. Remarkably, the shape of the spectrum remains the same, suggesting finite size scaling [43]. Finally, Fig 4D shows how the maximal dimensionality of the model *L* and the value of *k*_*c*_ depends on the network size *N*.

For small neural populations sizes, *k*_*c*_ *∼ L*, but as *N* increases, the number of possible interactions (L)grows, requiring longer recording lengths. Black circle denotes the network size used in retinal spike train data shown in this article (*N* = 50).

The fact that a larger number of neurons *N* requires longer recording lengths brings as consequence a value of *k*_*c*_ departing from *L*. As either the simulated or real data has a finite length *T*, increasing *N* generate an increasing number of unobserved monomials, or, monomials with very low occurrence probability, reducing the effective *χ* matrix rank. The effect of unobserved or very low empirical probability events on *χ* and, consequently, on *k*_*c*_ estimation is detailed in the next subsection.

### A measure of “code compressibility”

We now relate *k*_*c*_ to a notion of “neural code compressibility”. We start from an important remark: as a sum of covariance matrices, *χ* is non negative. Nevertheless, it can have many zero eigenvalues for two distinct reasons:

1. There are hidden linear dependencies between the coefficients *h*_*l*_. This is typically due to an hidden structure in the dynamics which has generated the data. An explicit example is provided in [13] where the *h*_*l*_ of a neuronal network model are computed as a function of the *W*_*ij*_s (synaptic weights) and stimulus. These eigenvalues are intrinsic to the dynamics and constitute somewhat the redundant part of the information that we want to remove to “explain” data. We note *d*_*N*_ the number of these eigenvalues.
2. *χ* is computed from finite rasters. Here, some monomials have zero empirical value when the corresponding event do not appear in the empirical raster. If *m*_*l*_ is one of these unobserved monomials we have *π*(*m*_*l*_) = 0 and, *∀l′*, *π*(*m*_*l*_*m*_*l ′*_) = 0, where *π* is empirical probability. We call this type of events Unobserved Events (*U*_*E*_). As a consequence, *χ*_*ll′*_ = 0, *∀l′* and the row *l* of *χ* has zero entries. Consequently, this row generates, in the spectrum of *χ*, a zero eigenvalue.

We have therefore ℛ + *d*_*N*_ + *U*_*E*_ = *L* where ℛ is the dimension of the image of *χ*. In general *k*_*c*_ *< R* because, after the cut-off there are eigenvalues, small, but nevertheless positive. If the cut-off is sharp, as we observe, *R* is very close to *k*_*c*_ though. On this basis we define the “compressibility” of the code as:

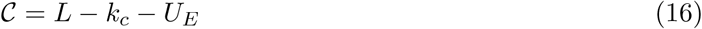

For sharp cut-off it is very close to *d*_*N*_ which precisely reflects the irrelevant dimensions in the space of parameters, corresponding to hidden dependencies.

### How do *k*_*c*_ and 𝒞 depend on the raster density

Using the aforementioned approach to dissect the relevant from the irrelevant dimensions, we aim to study the dependency between the raster density and code compressibility given a spatio-temporal MEM imposed to data. We consider neural density (or, conversely, sparsity) as the number of spikes of a raster divided by the number of bins of the raster. This density depends on many factors as the excitatory-inhibitory balance, network connectivity and stimuli, among others. Very dense neural responses could produce artificial neural correlations beyond the underlying statistics. On the opposite, very sparse responses could not provide enough information to fairly recover the correct empirical statistics of complex events (pairwise with time delays). Having this in mind we generated synthetic data with different levels of density and estimated *k*_*c*_ under these different regimes.

We generated different synthetic sets of neural data where the density of the response varies. To do this, starting from PWR2 model, we generated five new responses scaling the parameters of the MEM by a factor *β* = {0.4, 0.6, 0.8, 1.2, 1.4}^2^

The effect of increasing the parameter *β* has therefore a tendency to spread the distribution of parameters *h*_*l*_ as shown in Fig 5A enlarging its variance, where the parameters *h*_*l*_ associated to firing rates are represented in blue, and those associated to spatio-temporal interactions in red. As expected, scaling the model parameters by low values of *β* condense the parameters distribution.

**Figure 5:**
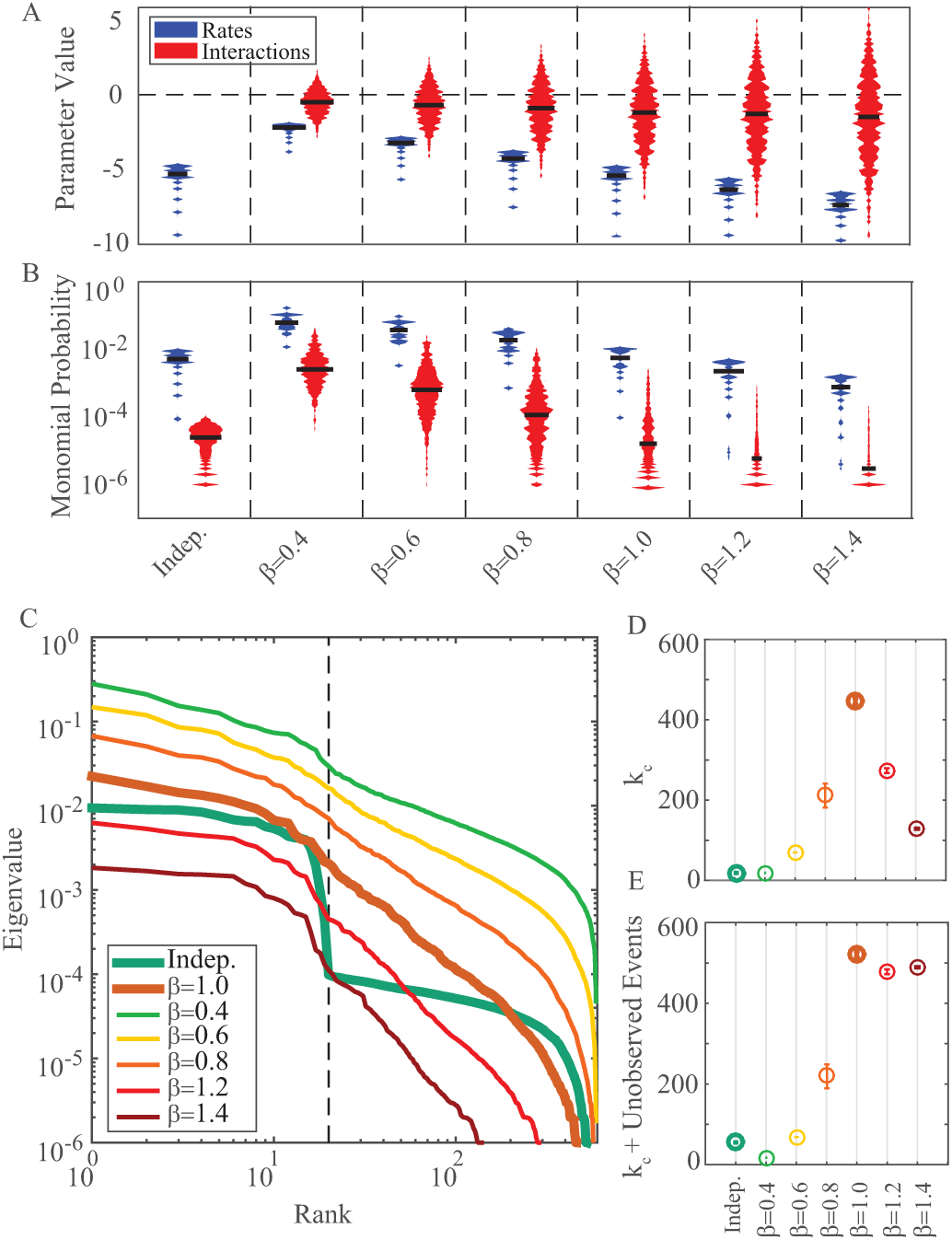
Dimensionality reduction on scaled PWR2 statistics. A: Underlying parameters distribution, split by firing rates (blue) and pairwise interactions (red). The bigger the scaling factor *β*, the more negative the rates parameters and the wider the interactions parameters distribution. *β* = 1 is the original PWR2 raster. The first column show the Independent raster as a reference. B: Corresponding monomials probabilities for the scaled rasters, split between firing rates and interactions, as in A. We see that increasing *β* has the effect of decreasing both rates and pairwise interactions probabilities, reaching the point where many pairwise interactions vanish (*β >* 1). C: Corresponding *χ* eigenvalue spectra (average out of 10 rasters for the scaled rasters). Increasing *β* flattens the spectrum, the spectrum offset and the number of eigenvalues above the minimal observed probability (1*/T*). D: *k*_*c*_ values for the scaled rasters. None of the scaled rasters shows *k*_*c*_ value as the one obtained for the original PWR2 raster. Dots represent the average of *k*_*c*_ over 10 different rasters with the same underlying parameters. Error bars represent 1 s.d. E: Average *k*_*c*_ values plus the number of unobserved events (*UE*). For *β >* 1 adding the unobserved events yields values close to the original PWR2 raster, showing that the dimensionality reduction obtained for those rasters is given mainly by the unobserved effects. For *β <* 1 we have less unobserved events, so the dimensionality reduction obtained for those cases is given mainly by the increased density of the raster.

Similarly, monomial probabilities are also affected by the *β* parameter (see Fig 5B). As expected, decreasing the value of *β <* 1 tends to equalize the probabilities of the monomials generating a very dense neural response. On the contrary, higher values of *β >* 1 tend to maximize the energy generating configurations of the response with less variability (sparse distribution). High values of *β* diminishes therefore the probability of occurrence of those monomials whose *h*_*l*_ value is negative while it increases the probability of monomials (spike events) with a positive *h*_*l*_.

Increasing *β* value causes two main effects on the eigenvalue spectrum of the *χ* matrix (see Fig 5C): (i) the spectrum is flatter as *β* decreases, (ii) there are more zero eigenvalues, i.e. *U*_*E*_ increases. Both effects come from the fact that the raster sparsity decreases with *β*, decreasing the events probabilities as well. Thus, this low probability events are reflected on the increase of *U*_*E*_ due to finite sampling and also on the flatness of the spectrum, given that the few observed events have similar probabilities. Decreasing the value of *β* below 1 decreases the value of *k*_*c*_. In a similar manner, values of *β* larger than 1 monotonically decrease the value of *k*_*c*_. Then, moving away from *β* = 1 reduces *k*_*c*_, but the reasons for the first case are different from the second. To illustrate this, Fig 5D shows the mean *k*_*c*_ value obtained for 10 different rasters (sharing the same underlying parameters and *β*) as a circle and 1 s.d as error bars. As we previously mentioned, small values of *β* produces a very dense neural response, that apparently can be condensed in a few dimensions (*k*_*c*_ = *N* for *β* = 0.4). On the second case, for *β >* 1 many events have very low probability, increasing *U*_*E*_, which will decrease *k*_*c*_. This difference is illustrated on figure 5E), where larger values of *β* generates responses with large *U*_*E*_.

For values of *β <* 1 we see almost no difference between Fig 5D and Fig 5E, meaning that *k*_*c*_ is an indicator of data compression not affected by the unobserved events (that are negligible). Nevertheless, the data compression detected by a low value of *k*_*c*_ is artificial because most of the correlations are induced by the effect of a high activation in the neural population. On the opposite, in the cases where *β >* 1 the unobserved events become significant and the low value of *k*_*c*_ obtained in Fig 5D was mainly due to the number of non-zero dimensions of the *χ* matrix without compression.

Thus, from this approach, the full dimensionality of the model imposed to data can be dissected on the effective dimensionality, *k*_*c*_, and the compressibility, 𝒞, reporting the level of redundancy that the model can capture from data.

### Dimensionality reduction on retina data

We are now interested in verifying if the properties found in synthetic data are scalable to real neural data. We did it on retina data obtained for three different conditions: photopic spontaneous activity (PSA), white noise (WN) and natural movie (NM)

Retina data was recorded in a 252-MEA system obtaining the response of 867 retinal ganglion cells in four different pieces of *O. degus* retina. Additionally, we generated shuffled version of the real data to compare the value of *k*_*c*_ and *C* in the two cases, where the shuffled version maintained the firing rates of each neuron destroying the spatio-temporal correlations. In Fig 6 we show the distribution of the empirical monomials present in real versus shuffled data. Monomials related to firing rates are shown in blue and those related to spatio-temporal correlations are shown in red. Interestingly, distribution of the pairwise interactions probabilities of the empirical recordings and their shuffled version for all stimuli are not significantly different (bin 10ms, Mann-Whitney test *P >* 0.05), so we don’t expect big differences on monomials empirical probabilities, but we do expect differences on the linear dependencies between them. Also from Fig 6 we observe that the highest activity, both for firing rates and spatio-temporal interactions, is obtained for NM, followed by WN and then by PSA. Thus, we modify both the raster density and the hidden dependencies of the neural activity by means of stimuli, where, in this case, the raster density increases with the stimuli high-order correlations.

**Figure 6:**
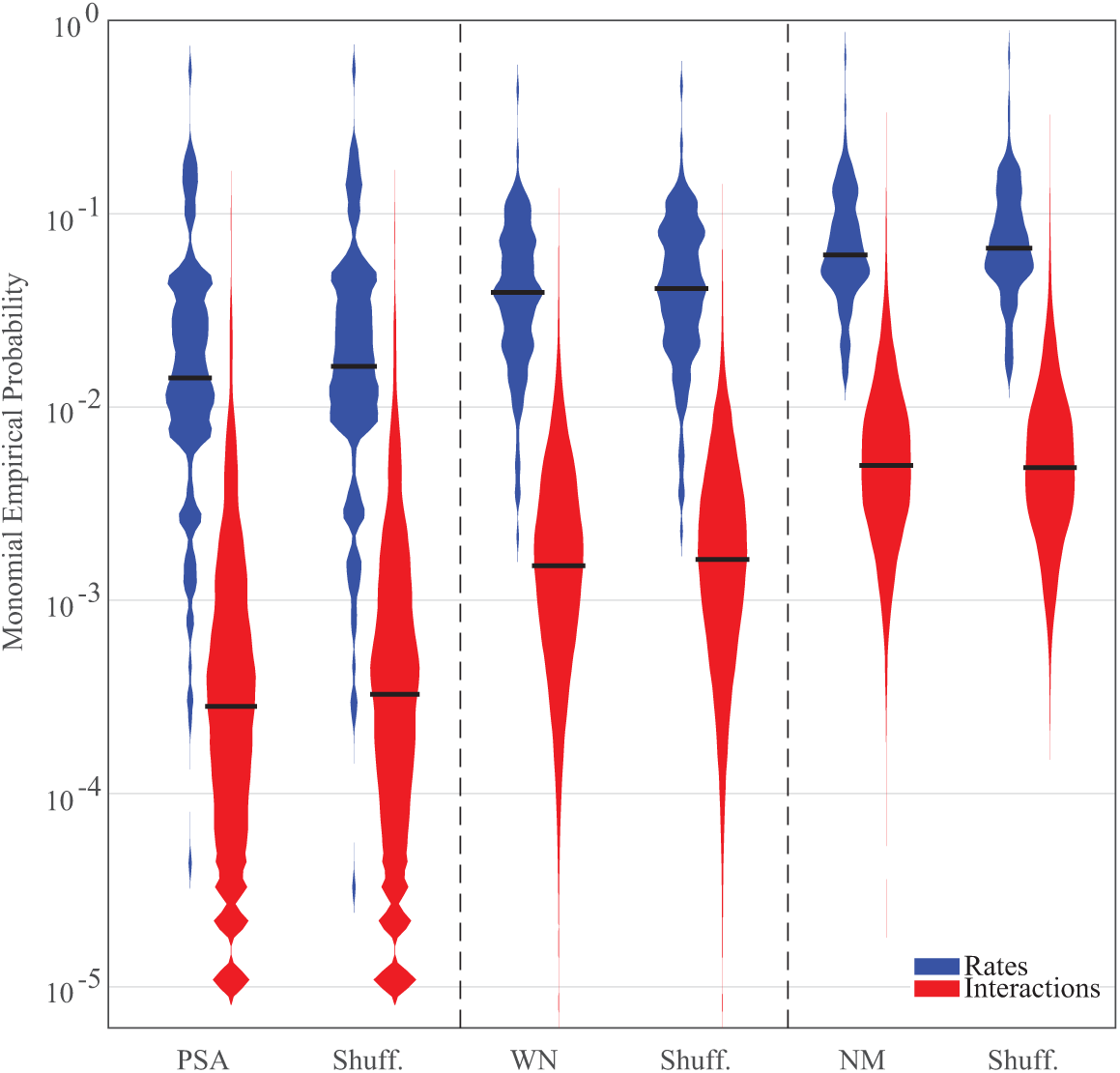
Distribution of firing rates and pairwise interactions. Violin plot of the firing rates (blue) and pairwise interactions (red) are shown for the 3 stimuli used and their corresponding shuffled version using a bin of 10ms (black horizontal line is the median). Both type of monomials increases with the stimuli high-order correlation. Note that the shuffled version reproduces the distribution of the pairwise interactions observed probabilities, even when the temporal structure of the raster is destroyed. This doesn’t imply that the linear dependencies between MEM parameters are kept.

### Experimental versus shuffled spectrum

We computed the *χ* eigenvalue spectrum of 30 random sub-networks (i.e. sub-samples of the entire population) of *N* = 50 from the total number of neurons recorded in each of the four experiments, under the 3 stimuli conditions. According to our synthetic rasters experiments, for *N* = 50 and *T ∼* 10^6^ we can get reliable *k*_*c*_ estimates (Fig 4D), which fits with our experimental recordings. Same procedure was applied to shuffled data.

Similar to what we obtained for scaled synthetic rasters (Fig 5C), we see that the spectrum is flatter, and the vanishing eigenvalues (below 1/T) of *χ* spectrum increases with the raster density, which is driven by the stimuli (Fig 7A). The size of the network (*N* = 50) is represented by a vertical dashed line. In real data, only WN condition shows a cut-off in the spectrum close to *N*. Interestingly, in shuffled data both NM and WN present this cut-off near *N*, suggesting that in the experimental recordings there are significant linear dependences between monomials that are not present in the shuffled version (see Fig 7B). Specially, NM shows a smooth decay of the spectrum, similar to the observed for PWR2, which is highly modified when the raster is shuffled, having a sharp cut-off close to *N*. So, even when the distribution of the probabilities of firing rates and pairwise interaction remain the same after the shuffling procedure (Fig 6), the linear dependencies between them are modified, changing the shape of the spectrum, showing sharper cut-offs for all conditions.

**Figure 7:**
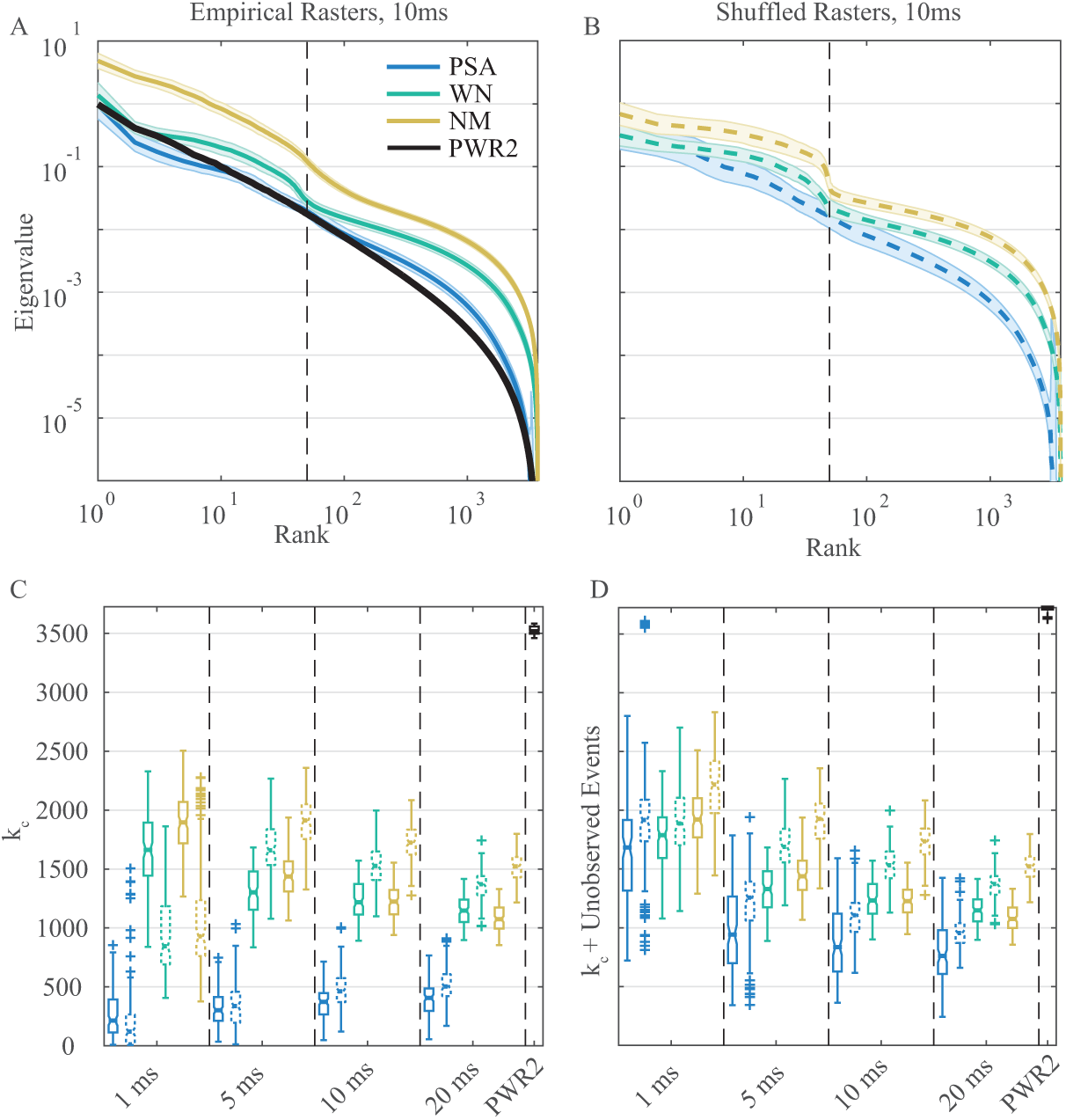
Dimensionality reduction on RGC data. A: Average spectrum (30 random sub networks plus 1 s.d., solid line and shaded area, respectively) of RGC data (*N* = 50) under 3 different stimuli conditions, with 10ms bin size. Stimuli high-order statistics increases the spectra offset (the magnitude of the eigenvalues). Except for WN, there is no clear cut-off close to *N*. Black line is a random PWR2 raster of the same network size than the RGC raster plot. B: Same than A, but for the shuffled version of the empirical rasters in dashed lines. All of them show a clear cut-off close to *N* and, preserving the effects induced by stimuli high-order correlations, i.e. change on the first eigenvalue and offset. C: Box plots for *k*_*c*_ values for empirical (solid) and shuffled rasters (dashed). Given the absence of prior knowledge about the relevant timescales the brain uses to integrate retinal signals, we studied several bin sizes. For example, for 1ms, *k*_*c*_ is higher for shuffled data in all bin sizes, under all conditions. Stimuli high-order correlations significantly increases *k*_*c*_. Also, neither of both type of rasters reaches the *k*_*c*_ values obtained for a PWR2 rasters, which shows almost no dimensionality reduction. For fast time scales (1 and 5ms), *k*_*c*_ increases with the stimuli-high order correlations, but at 10ms WN and NM are not significantly different and for 20 ms WN is larger than NM. D: Same as C, but corrected by the number of unobserved events. Now for all bin sizes the shuffled data has bigger values than the empirical one. Shuffled data has values almost a half than a PWR2 raster of the same size, regardless of the absence of linear dependences by construction. The effect of stimuli high-order correlations on the effective dimensionality remains.

### Computing *k*_*c*_ for retinal ganglion cells spike trains

We computed *k*_*c*_ for spike trains of retinal ganglion cells obtained for different stimulus. For each stimulus we generate different spike trains (experimental and shuffled) ans we use different time bins to binarize the data: 1, 5, 10 and 20ms. As a global picture, the value of *k*_*c*_ in the real and shuffled data increases as the activity of the network does (Fig 7C). This behavior is replicated for all the bin sizes.

The *k*_*c*_ analysis on retinal data reveals that the RGC activity is not random, showing almost the half of dimensionality compared to a random PWR2 with the same network size, which is shown in black in the upper right corner of Fig 7C. Furthermore, even if the firing rates and spatio-temporal events have the same distribution between real and shuffled data, the values of *k*_*c*_ obtained in the two cases differ, the real value being always smaller than the shuffled version. For all the time scales (from 1 up to 20 ms) we observe that the value of *k*_*c*_ in the shuffled data is always higher than the one obtained for real data. Additionally, for the shuffled data the value of *k*_*c*_ monotonically increases with the activity. Similarly, the real data also presents a tendency to increase the value of *k*_*c*_ as the raster density does; nevertheless the relation is inverted between WN and NM for bin sizes 10 and 20ms: in these two cases the *k*_*c*_ value obtained for NM is smaller than the values obtained for WN.

In order to verify if the *k*_*c*_ values are actually associated to linear dependencies between neurons, and not to unobserved event, we did the correction *k*_*c*_ + *U*_*E*_ as it was shown in Fig 7D. PSA has the larger *U*_*E*_ values, showing a big difference between *k*_*c*_ and *k*_*c*_ + *U*_*E*_. In general, *k*_*c*_ corrected by *U*_*E*_ decreases with the bin size, because the larger the bin, the more likely is to observe two spikes on the same bin and less likely is to haver unobserved events. As a consequence, this increases the apparent interdependence of RGC population activity (both for real and shuffled data). Specifically, we see that *k*_*c*_ increases with the stimuli high-order correlation for fast time scales (Mann-Whitney test, *P <* 10^*-*5^ for all comparisons). This suggests that for fast time scales the retina increases the number of coding channels as the stimuli high-order correlation increases. However, for medium time scales the pictures changes, showing no significant differences between WN and NM for 10ms (Mann-Whitney test, *P >* 0.1) and showing higher *k*_*c*_ values for WN than NM for 20ms (Mann-Whitney test, *P <* 10^*-*5^). This suggest that at larger time scales the RGC activity under NM becomes more interdependent than for WN, which could be related to the increased level of redundancy of NM compared to WN and the time scale that this redundancies are captured by the retina generating a redundant activity.

On the other hand, the shuffled data show higher *k*_*c*_ values for almost all the bin sizes and conditions (Mann-Whitney test, *P <* 10^*-*4^), except for 1ms where is significantly small (Mann-Whitney test, *P <* 10^*-*4^) and for PSA at 5ms, where they are not significantly different (Mann-Whitney test, *P >* 0.1). However, when corrected by the number of unobserved events (Fig 7D), the picture is the same for all bin sizes: shuffled data have always higher *k*_*c*_ values than the real ones (Mann-Whitney test, *P <* 10^*-*6^ and *P <* 0.01 for WN at 1ms). This confirms that the RGC neural code has interdependences that can be mapped onto a lower dimensional space (i.e. compressed), compared to the shuffled version, that lacks of interdependences by construction. Yet, the *k*_*c*_ obtained for the shuffled raster is still very small (almost a half) compared to the random PWR2 raster, which suggests that just the firing rates distribution, i.e. the diversity of firing rates, introduces some non-random interdependences in the neural code, allowing compression. Finally, for medium time scales (10-20ms), shuffled data shows an increase of *k*_*c*_ (an the correction by *U*_*E*_) with the stimuli high-order correlation, on the opposite to empirical data, where at this time scales NM induces a lower dimensionality, compared to WN. This support the idea that there are time scales where the stimuli redundancies are reflected on the retinal activity and that this timescale is not present anymore on shuffled data.

In sum, the density of the RGC rasters increases with the stimuli high-order correlations. Also, RGC neural code is highly compressible compared to a random raster of same network size. Even compared to shuffled data, the RGC neural code is more compressible. Furthermore, a significant compression can be achieved considering just the firing rates distribution of empirical data, suggesting an effect of the diversity of firing rates on the code compressibility. Although stimuli high-order correlations increases the raster density, this activity can be compressed on a lower dimension for NM than WN using a bin of 20ms. Thus, an increased raster density induced by stimuli does not imply necessarily more coding channels. Instead, an increased raster density could be mapped onto a low-dimensional hidden structure, as in the case of concomitance of dense activity and oscillations. Regarding the time scales, *k*_*c*_ + *U*_*E*_ is inversely proportional to the bin size for all stimuli. On the one hand, this could be due to the specific time scales at which more redundancy is present on the neural activity. On the other hand, this could be due to artifactual correlations induced by binning. Both scenarios are possible and not mutually excluding, however, the problem of binning neural data is far from being solved.

## Discussion

In this paper we have proposed a method to reduce the dimensionality of MEM on artificial and biological spiking networks. It is grounded on information geometry via the matrix *χ*, which characterizes how a small variation of parameters impacts the statistical estimations. The *χ* matrix captures the interdependences between the neural code variables. After an eigen-decomposition process, the eigenvalue spectrum of *χ* exhibit two cut-offs. The first one shows that, both in synthetic as well as in retina data, a large part is “explained” by the firing rates of the neurons. Conversely, the second cut-off (here called *k*_*c*_) reflects a non trivial effect associated with higher order statistics. As the eigendirections on the right part of the second cut-off correspond to noise, the spectrum lying between the two cut-off contains a relevant information associated to statistics of higher order.

The reduction of the MEM dimensionality is directly linked with data compression, obtained from the linear dependencies between variables of the MEM that reflects hidden (non linear) interactions between neurons. For example, our analysis in the case of synthetic rasters, where in contrast to retinal data both the firing rates and the pairwise interactions are defined randomly, shows a *k*_*c*_ value very close to *L* (maximal dimensionality). This demonstrates that if there are no linear relationships between the parameters by construction, the code is not compressible and we have almost one dimension per parameter. On the opposite, retina data shows a significant compression of *∼* 50%, as expected from a neural tissue where cells are driven by common inputs and cells are electrically coupled [26], increasing the level of dependency between them. This compression reduces the dimensionality of the MEM to a lower dimensional space, where each dimension is a linear combination of model parameters, characterizing the population activity by a set of independent dimensions representing the inner structure of the network activity.

### Limits of the method

The first limitation of our method comes from the numerical approximation used to compute *χ* matrix, which is obtained by summing monomials correlations at different time lags (Eq. 11). An exact exponential decay of correlation function with time will ensure the convergence of the series. But, because we are estimating correlations from finite rasters, it is hard to estimate correlations with large time lags and errors accumulate.

To truncate this approximation we need to consider a trade-off between temporal resolution and reduction of noise. To this end we use 4 time lags (i.e. 4 terms of the sum), which is equivalent to the double of memory depth used in the model (*R* = 2). This numerical estimation of *χ* also imposes limits on the method, given that considering too large *R*s will generate a *χ* matrix that is governed by noise.

On the other hand, it is possible to compute *χ* from the model parameters, requiring a previous model fitting step, as done by [25]. However, for *N >* 20 and *R >* 1 the MEM computation becomes prohibitive as the network size and the memory depth of the model increases. So, despite the numerical approximation and its intrinsic errors, computing *χ* from the empirical monomials time correlations is the best approach we found to work with medium size networks and spatio-temporal constrains.

Nevertheless, without fitting the MEM model we miss information about the sign (positive or negative) and the magnitude of the interactions (weak or strong) defining the the network topology and statistics. However, the scope of this work was not to fit different models on data and test its performance (e.g. Bayesian or Akaike information criterion that takes into account the model parameters and likelihood [27]) neither study changes in network topology under different stimuli. Instead, we focus on exploring the geometrical properties of the MEM and its meaning in terms of the neural code redundancy and compressibility. As we presented here, these geometrical properties can be directly extracted from the *χ* matrix without fitting a MEM.

### *k*_*c*_ **is not one value, but a set of values**

The main challenge this methodology introduces is the selection of *k*_*c*_. The selection of *k*_*c*_ is related to the minimization of the so-called confidence volume, which not only depends on the number of dimensions given by *k*_*c*_, but also on the imposed tolerance ϵ. So, strictly speaking, the selection of *k*_*c*_ depends on the tolerance ϵ that the analyst decide to use based on equation 22. However, here we used a criterion based on a trade-off between dimensionality and the minimal confidence volume (log *V*(*k*_*c*_, *E*)): as shown on Fig 2, log 𝒱(*k*_*c*_, *E*) decays monotonically as *k*_*c*_ increases and ϵ decreases, reaching an inflection point where even if we decrease the accuracy by orders of magnitude *k*_*c*_ remains almost unchanged. This means that we look for the minimal dimensions with the highest possible accuracy. Thus, our choice of *k*_*c*_ was not based on fixing a tolerance, but rather on relationship between the minimal confidence volume, the dimensionality and imposed accuracy.

### Comparison with similar analysis on spike trains

To our knowledge, there is no previous work related to the analysis of the *χ* matrix considering *spatio-temporal* pairwise interactions applied to neuronal networks.

The authors in [25] proposed a similar analysis considering only spatial interactions, i.e. the Fisher Information Matrix (FIM) for the Ising model on *in vitro*, *in vivo* and *in silico* networks. The work of [25] studies small neural networks (10 neurons), they looked for stiff neurons, which are related to stiff dimensions (the largest FIM eigenvalues), proposing that those neurons are the ones giving stability to the network, while the neurons involved on the sloppy dimensions (dimensions where the parameters can have significant changes without affecting the model) are the ones involved on plasticity, allowing the network to remodel its connections.

Similarly, using larger network sizes (*N* = 50) and spatio-temporal interactions, our approach exploits the linear dependencies of neuron interactions (given by the *χ* matrix) to find a minimal set of dimensions that better represents the neural code, which could be considered an equivalent to stiff dimensions present in [25].

To this end, we found two set of stiff dimensions: the ones before the first cut-off, related mainly to neuron firing rates and the second set, after the first cut-off, related to spatio-temporal interactions. According to our framework, the sloppy dimension would be the ones beyond *k*_*c*_, but we could also interpret the first *N* dimensions as the stiff dimensions, the ones between *N* and *k*_*c*_ as the sloppy dimensions and the ones beyond *k*_*c*_ just noisy dimensions. This re-interpretation arises from the large magnitude difference between the first and second set of dimensions and is compatible with our previous description of compressibility: to represent data with the minimal set of dimensions we require both stiff and sloppy dimensions. So, we consider the sloppy dimensions as relevant, because we need them to “explain” data, while we consider the irrelevant dimensions as noise.

Recently, Battaglia et al [28], studying large scale networks between brain areas, proposed a concept called *Meta Connectivity*, which instead of analyzing the correlation between nodes of a network (the usual functional/effective connectivity analysis), focuses on the correlations between the network interaction along time. This means focusing not on the coupling *ω*_*i*_(0)*ω*_*j*_(0) between neurons *ω*_*i*_ and *ω*_*j*_, but focusing on the interactions between the couplings *ω*_*i*_(0)*ω*_*j*_(0) and *ω*_*k*_(0)*ω*_*l*_(0). This provides information about instantaneous high-order correlation for at least 3 nodes of the network (e.g. case of *i* = *k*) and captures the relationships between modules of network activity. Thus, *χ* matrix is both a functional/effective connectivity matrix (the matrix entries related to the correlations between firing rates) and also a meta connectivity matrix (the matrix entries related to correlations between pair-wise interactions), which also includes information about the temporal interactions between network nodes and network modules. The extension of our analysis towards the understanding of the meta connectivity has never been applied to networks of neurons. It is a future research direction, where the focus is on the variability and dependence of the interactions of a network.

### Stimuli-induced changes on RGC population activity

Retina data has significant high-order correlations, including pairwise spatial[2, 3], temporal [16] interactions, triplets [9] and groups of neurons [10]. These correlations have been widely studied under MEM, which can accurately reproduce the raster spatio-temporal patterns. Nevertheless, there has been no work devoted to reduce the MEM dimensionality considering the inner dependences between the population activity variables.

Here, as a proof of concept, we used retina data of a diurnal rodent under 3 different stimuli with different statistics, from photopic spontaneous activity (spatio-temporal uniform full field, no second order statistics), a spatio-temporal white noise (gaussian statistics) to a repeated natural movie (high-order correlations both on time and space). From our analysis, we know that these stimuli high-order correlations increases the magnitude of both the firing rates and the raster high-order correlations, even making silent cells to fire. This could be related to recruitment of specific cell types by the stimuli features (e.g. local contrast [29], optic flow, color [30], among others).

In general, most of the correlated activity observed in the retina could be attributed to the receptive field overlap between recorded RGCs and to shared common noise [31]. Additionally, electrical coupling are highly present in the *O. degus* retina [26] inducing fast correlations between close cells. Also, there could be correlations driven by amacrine cells, but the mechanisms and roles of those cells on the *O. degus* retina is still unknown. The aforementioned causes of correlations modifies the pairwise spatio-temporal interactions observed in real data. Furthermore, we explored the presence of these correlated activity in the neural code. To do this we compared the results of recorded data with a shuffled version of it, which preserves the firing rates distribution. Interestingly, both data sets share the pairwise interactions probability distribution suggesting, in this case, that second order statistics can be preserved just fixing the firing rates distribution.

Finally, as a main conclusion, our method suggest that RGC activity has significant high-order statistics that are modified by stimuli, compared to shuffled data. Thus, this significant interactions on RGC data are the base of the increased compressibility compare to shuffled data.

### Compressibility of the RGC code

In order to study the compressibility of the RGC code, we studied *χ* spectrum and *k*_*c*_ for RGC under three stimuli conditions, finding that RGC population code adapts to stimuli conditions by changing the number of independent channels.

The first difference we found between stimuli was on *χ* spectrum, which shows an increase on the offset, i.e. the eigenvalues increase their magnitude as the stimuli high-order correlation increases. We recall that the stimuli-high order correlation increases the raster density. But given the way *χ* is computed (see Eq. 11), the shape of the spectrum comes not only from the increased monomials probabilities, but also from the dependence between the set of MEM monomials, all of them captured by the matrix. On the one hand, shuffled data also exhibit these differences on the eigenvalue magnitudes (Fig 7B), showing that eigenvalues magnitude are closely related to the raster density (also shown for synthetic data on Fig 5C). On the other hand, the differences between the cut-off for empirical and shuffled data would be related to the linear dependences between the monomials, that in the latter case are destroyed. Then, using shuffled data as a control, we suggest that the magnitude of the eigenvalue spectrum highly depends on the monomials probability (raster density) while the cut-off depends on the hidden linear dependencies between them.

The second difference we found between stimuli is on *k*_*c*_ and 𝒞 i.e. our approximation to the compressibility of the neural code. As expected from the stimuli statistics and retinal stimuli integration, *k*_*c*_ and 𝒞, are always lower for PSA than for the dynamic stimuli. This suggests that the network optimizes the number of dimensions required for coding the stimuli depending on the stimuli statistics. In terms of metabolic cost, a very redundant stimulus as PSA (which has the same spatial and temporal information all over the stimuli space), may be coded with less dimensions than stimuli with more independent components (less redundant), thus, optimizing the metabolic resources.

However, for bin sizes of 1 and 5ms we observe that *k*_*c*_ is higher for NM than for WN. This relation varies for larger values of bin sizes, such as 10 or 20ms. For bin sizes of 10ms WN has the same number of relevant dimensions than NM, while for 20ms WN has more dimensions than NM. So, for fast time scales, we face a non-optimal situation, because NM has more redundancies than the WN. It could be possible that at this time scales we are not capturing the inter-dependences of the neural code that are relevant for the brain. For example, at 20 ms bin size, we see the expected optimal effect: the system exploits the stimuli redundancies and exhibit more compression for NM than for WN. Coincidently, many MEM on retina have been done using 20 ms as bin size [2, 3], which in our case is the bin that allows the highest compression. In addition, synthetic data shows that if the raster is too dense, the underlying statistics hides under the noisy activity, which could also be possible at large values of bin sizes. To control this situation we used shuffled rasters, which preserves the same raster density. In the shuffled rasters we see that *k*_*c*_ increases with the raster density, discarding the effect of density on the changes of dimensionality at higher bin sizes.

Thus, at large bin sizes, the interdependences of the neural code are responsible for the compression effect and not the raster density hiding some events. Nevertheless, we do not know in advance what time scale(s) is(are) actually relevant for the brain and neither if our assumptions about code optimality and the neural code variables (firing rate and spatio-temporal interactions) are right, so the choice of the bin size and code variables is still an open question and somewhat arbitrary.

Finally, our work is related to the idea that a stimuli-dependent network noisy spiking neurons adapt its code according to noise and stimuli correlations [21], instead of using just one way of coding. In our case, the stimuli-dependent network is a biological one, so we don’t have access to modify the noise of each neuron nor the network noise. Instead, we can just modify the stimuli correlations, which changes the dimensionality of the code. This change in dimensionality could reflect the smooth interpolation between encoding strategies: highly redundant stimuli evokes fewer dimensions than stimuli which presents high-order correlations.

This suggests that the MEM dimensionality is also a measure of the code redundancies. The analytic relationship between dimensionality reduction and the coding strategies require an extensive mathematical and computational research that is not developed here, however, we provide an indirect way of studying the interdependences of the neural code as the stimuli conditions changes.

## Supplementary Material

**Gibbs measures in the sense of Bowen.** Consider a potential *H*_*h*_ of range *R ≥* 2. A shift invariant probability measure *μ* is called a Gibbs measure (in the sense of Bowen) if there are constants ℳ *>* 1 and 𝒫[ ℋ_*h*_] ∈ ℝ s.t.

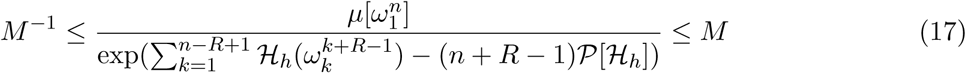

It is easy to see that the classical Boltzmann-Gibbs distribution is a particular case of (17), when ℳ = 1 and ℋ is a potential of range 1.

**Ruelle-Follmer theorem**: Suppose *μ′* is a Gibbs measure for some potential ℋ _*h*^*′*^_, and *μ* is another Gibbs measure. Then the relative entropy density:

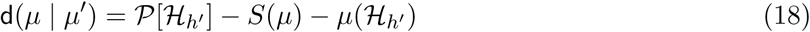

if d(μ | μ ′) = 0, we obtain the variational characterization of Gibbs measures (4).

Following [36], consider the potential 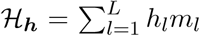 associated with an ergodic Markov Chain *μ*(*P, π*). Consider a sample of *μ*(*P, π*) of length *n* and the observables 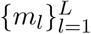We may obtain from the sample new maximum entropy parameters ***h′***. The probability that the maximum entropy parameters ***h′*** associated with an ergodic Markov Chain *μ′*(*P ′, π′*) get close to ℋ follow the asymptotic relationship:

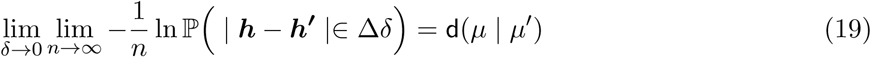

where Δ*δ* = [*-δ, δ*]^*K*^. Choosing Δ*δ* close to 0 we may formally rewrite the above relationship in the form:

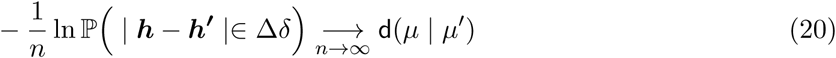

Thus, for large *n*,

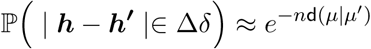

Meaning that close-by parameters index very similar distributions [41].

Consider two maximum entropy Markov chains *μ*(*P, π*) and *μ′*(*P ′, π′*) specified by *H***_*h*_** and *ℋ*_*h′*_ respectively. As both satisfy the variational principle, we have that the relative entropy (18) reads:

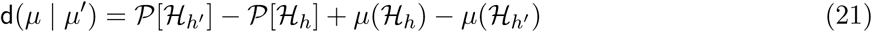

Taking the expansion of d(*μ | μ′*) around ***h′*** = ***h*** we obtain:

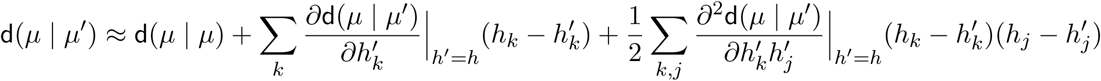

Since d(*μ | μ′*) is minimized at *h′* = *h*:

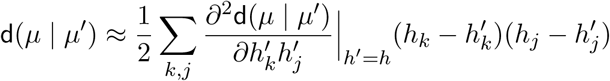

Taking the second derivative of d(*μ | μ′*) from (21), we obtain:

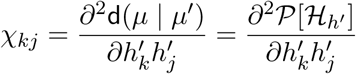

Given two maximum entropy Markov chains specified by *H***_*h*_** and *ℋ*_*h′*_ in the limit of large *T* they are *E*-indistinguishable if:

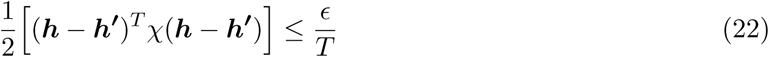

where *χ* is the Fisher information matrix, which represents the curvature of the relative entropy.

## Materials and Methods

### Ethics Statement

Animal manipulation and breeding and corresponding experiments were approved by the bioethics committee of the Universidad de Valparaiso, in accordance with the bioethics regulation of the Chilean Research Council (CONICYT) and international protocols.

### Animals and Recordings

4 Adult male and female Octodon degus (3-6 months) were maintained in the animal facility of the Universidad de Valparaiso, at 20–25^*º*^C on a 12-h light-dark cycle, with access to food and water adlibitum. The methods of MEA recording has previously been described [26]. In brief, animals were euthanized under deep isofluorano or halothane anesthesia and both eyes were extracted. Then, one of the extracted retinas was diced into quarters while the other was stored in oxygenated in oxygenated (*O*_2_ 95% *CO*_2_ 5%) AMES medium at 32^*º*^C in the dark for further experiments. The same AMES media was used for continuous perfusion during extracellular recordings. For MEA recordings (MEA USB-256, 20kHz sample, Multichannel Systems GmbH, Germany), one piece of retina was mounted onto a dialysis membrane placed into a ring device mounted in a traveling (up/down) cylinder, which was moved to contact the electrode surface of the MEA recording array. Data were processed off-line using Plexon Offline Sorter (Plexon Instruments). Further, spikes were detected using a threshold of −4.5 to −5.5 S.D. from the mean voltage value and then were manually classified using the 2D space of the first two principal components on each electrode. Only somatic spikes were kept. Refractory period violations were detected and discarded if two or more spikes of the same neuron occur in a 2ms period. We recorded on 3 stimuli conditions (see next section): i) Photopic Spontaneous Activity (PSA), ii) Spatio-temporal white Noise (WN) and iii) Natural Movie (NM), obtaining 151, 200, 246 and 270 RGC for each of the 4 experiments, respectively. For each experiment 30 random subsamples of 50 neurons were taken and *χ* matrix corresponding to a Pairwise R=2 model were computed considering 4 bin sizes (1, 5, 10 and 20 ms) for each subsample, yielding 30 *k*_*c*_ values per experiment, per condition and per bin size. The shuffled version of these rasters were submitted to the same analysis.

### Visual Stimuli

Visual stimuli were generated by a custom software created with PsychoToolbox (Matlab) on a Mini-Mac Apple computer and projected onto the retina with a LED projector (PLED-W500, Viewsonic, USA) equipped with an electronic shutter (Vincent Associates, Rochester, USA) and connected to an inverted microscope (Lens 4x, Eclipse TE2000, NIKON, Japan). The image was by 380 × 380 pixels, each covering 5*μm*^2^. Since rodents are dichromatic (green and blue/UV cones), in our experiments only the B (blue) and G (green) beams of the projector were used, while the R (red) channel was used for signal synchronization. Dark spontaneous activity was recorded in order to monitor the stabilization of the activity. The stimuli where applied. For PSA a space-time invariant stimuli with G and B intensities equal to the mean intensity of the NM stimulus were presented for 15 mins. WN stimulus with a block size of 50*μm* was used at a rate of 60 fps and presented for 20 mins, with each block taking independently 0 or 255 (max value) in the pixel value scale. NM consisted of an 1800 frames movie recorded on the natural habitat of the rodent using a robotic solution to capture the natural visual environment of degus, including grass, trees, optic flow, head-like movements. This short movie was presented 40 times at a refresh rate of 60fps, yielding a total duration of 20 mins. Optical density filters in the optical path were used to control final light intensity. A CCD camera (Pixelfly, PCO, USA) attached to the microscope was used for online visualization and calibration of the light stimuli projected onto the recording array.

### Generation of Synthetic Data

Synthetic rasters (*T* = 2.10^6^ time-points, *N* = 20 neurons) were generated using different underlying statistics: **Independent**, where only firing rates are defined, *L* = *N*; **Pairwise R=2** (PWR2), with firing rates and spatio-temporal correlations, *L* = *N* (3*N -* 1)*/*2. Underlying coefficients related to firing rates and to pairwise interactions were randomly chosen from a normal distribution with mean −5 and −1, respectively, and 1 standard deviation. were generated scaling the magnitude of the model parameters by a factor *β* = [0.4 0.6 0.8 1.2 1.4]. In addition, 6 more random PWR2 rasters were generated with *N* = [30 40 50 60 70 80] to study the dependence between *k*_*c*_estimation and the network size. Each raster was generated using the PRANAS software (https://pranas.inria.fr/) [45]. For the first 3 rasters, 100 random subsamples with half duration of the whole recording were taken for each raster and the *χ* matrix associated with a Pairwise R=2 model was computed. Then, *k*_*c*_ (see text) was found by volume minimization, yielding 100 *k*_*c*_ values per raster. For the scales rasters, the same procedure was applied, but using 10 temporal subsamples.

### Shuffling

In order to destroy the dependencies between the empirical raster monomials, we have generated random rasters where the number of neurons and firing rates was exactly the same than observed on the recordings (i.e. on each retina under each stimuli condition), but the spikes times were taken uniformly at random, avoiding violations of the refractory period (2ms) [44]

## Acknowledgments

Financial support: CONICYT-FONDECYT 1140403 and 1150638, CONICYT-Basal Project FB0008 (Chile), CONICYT-PAI Insercíon 79160120; Grant ICM-P09-022-F supported by the Millenium Scientific Initiative of the Ministerio de Economia, Desarrollo y Turismo (Chile); ECOS-Conicyt C13E06; ONR Research Grant #N62909-14-1-N121, ANR TRAJECTORY CE37 (France). We thank Felipe Olivares and Michael Pizarro to help in the experiments.

## Footnotes

1 Christoffel coefficients can be computed from the metric *χ*, giving thus access to the local information geometry and higher order statistics [24]

2 Increasing *β*, known in physics as inverse temperature, amounts to favor configurations maximizing the energy, i.e., monomials with high positive terms. On the opposite, decreasing *β* (increasing temperature) tends to equalize probabilities of patterns. Note that, in contrast to statistical physics, energy is maximized, not minimized, because we don’t have the minus sign in front of the potential, in the Gibbs distribution.

